# A missense variant in Mitochondrial Amidoxime Reducing Component 1 gene and protection against liver disease

**DOI:** 10.1101/594523

**Authors:** Connor A. Emdin, Mary Haas, Amit V. Khera, Krishna Aragam, Mark Chaffin, Lan Jiang, Wei-Qi Wei, Qiping Feng, Juha Karjalainen, Aki Havulinna, Tuomo Kiiskinen, Alexander Bick, Diego Ardissino, James G. Wilson, Heribert Schunkert, Ruth McPherson, Hugh Watkins, Roberto Elosua, Matthew J Bown, Nilesh J Samani, Usman Baber, Jeanette Erdmann, Namrata Gupta, John Danesh, Danish Saleheen, Mark Daly, Joshua Denny, Stacey Gabriel, Sekar Kathiresan

**Affiliations:** Center for Genomic Medicine, Massachusetts General Hospital, Boston, MA 02114; Department of Medicine, Harvard Medical School, Boston, MA 02114; Program in Medical and Population Genetics, Broad Institute, Cambridge, MA, 02142; Departments of Biomedical Informatics, Vanderbilt University, Vanderbilt, TN 3724; Departments of Medicine, Vanderbilt University, Vanderbilt, TN 37240; Institute for Molecular Medicine Finland (FIMM), University of Helsinki, FI-00014, Helsinki, Finland; Division of Cardiology, Azienda Ospedaliero–Universitaria di Parma, Parma, Italy 43121; Associazione per lo Studio Della Trombosi in Cardiologia, Pavia, Italy 27100; Department of Physiology and Biophysics, University of Mississippi Medical Center, Jackson, MS 39216; Deutsches Herzzentrum München, Technische Universität München, Deutsches Zentrum für Herz-Kreislauf-Forschung, München, Germany 80333; University of Ottawa Heart Institute, Ottawa, Ontario, Canada K1Y4W7; Division of Cardiovascular Medicine, Radcliffe Department of Medicine, University of Oxford, Oxford, United Kingdom OX12JD; Wellcome Trust Centre for Human Genetics, University of Oxford, Oxford, United Kingdom OX12JD; Cardiovascular Epidemiology and Genetics, Hospital del Mar Research Institute, Barcelona, Spain 08003; CIBER Enfermedades Cardiovasculares (CIBERCV), Barcelona, Spain 28029; Facultat de Medicina, Universitat de Vic-Central de Cataluña, Vic, Spain 08500; Department of Cardiovascular Sciences, University of Leicester, and NIHR Leicester Biomedical Research Centre, Leicester, United Kingdom LE17RH; The Zena and Michael A. Wiener Cardiovascular Institute, Icahn School of Medicine at Mount Sinai, New York, New York 10029; Institute for Cardiogenetics, University of Lübeck, Lübeck, Germany 23562; DZHK (German Research Centre for Cardiovascular Research), partner site Hamburg/Lübeck/Kiel, 23562 Lübeck, Germany; Department of Biostatistics and Epidemiology, Perelman School of Medicine, University of Pennsylvania, Philadelphia, PA 19104; Center for Non-Communicable Diseases, Karachi, Pakistan; Cardiovascular Epidemiology Unit, Department of Public Health and Primary Care, University of Cambridge, Cambridge, UK; Wellcome Trust Sanger Institute, Hinxton, Cambridge, CB10 1SA, UK; National Institute of Health Research Blood and Transplant; Research Unit in Donor Health and Genomics, University of Cambridge

## Abstract

Analyzing 5770 all-cause cirrhosis cases and 572,850 controls from seven cohorts, we identify a missense variant in the Mitochondrial Amidoxime Reducing Component 1 gene (*MARC1* p.A165T) that associates with protection from all-cause cirrhosis (OR 0.88, p=2.1*10^−8^). This same variant also associates with lower levels of hepatic fat on computed tomographic imaging and lower odds of physician-diagnosed fatty liver as well as lower blood levels of alanine transaminase (−0.012 SD, 1.4*10^−8^), alkaline phosphatase (−0.019 SD, 6.6*10^−9^), total cholesterol (−0.037 SD, p=1*10^−18^) and LDL cholesterol (−0.035 SD, p=7.3*10^−16^). Carriers of rare protein-truncating variants in *MARC1* had lower liver enzyme levels, cholesterol levels, and reduced odds of liver disease (OR 0.19, p= 0.04) suggesting that deficiency of the MARC1 enzyme protects against cirrhosis.

---

Cirrhosis is often considered to be the final stage of distinct pathogenic processes including excess alcohol consumption, fatty liver secondary to obesity and viral infection.^1^ However, analyses of these separate processes have identified similar genetic determinants. For example, *PNPLA3* p.I48M and *TM6SF2* p.E40K, although initially identified as associated with hepatic steatosis^2,3^, strongly predispose to the development of alcoholic cirrhosis^4^, non-alcoholic cirrhosis^5,6^ and hepatitis C-related cirrhosis^7,8^. The recently identified splice variant rs72613567 in *HSD17B13* similarly protects against alcoholic cirrhosis, non-alcoholic cirrhosis and severe liver fibrosis among individuals with hepatitis C (**Supp. Figure 1**).^9,10^ These individual findings suggest that some genetic variants may predispose to all-cause cirrhosis through pathways common to alcoholic and non-alcoholic cirrhosis.

**Figure 1.**
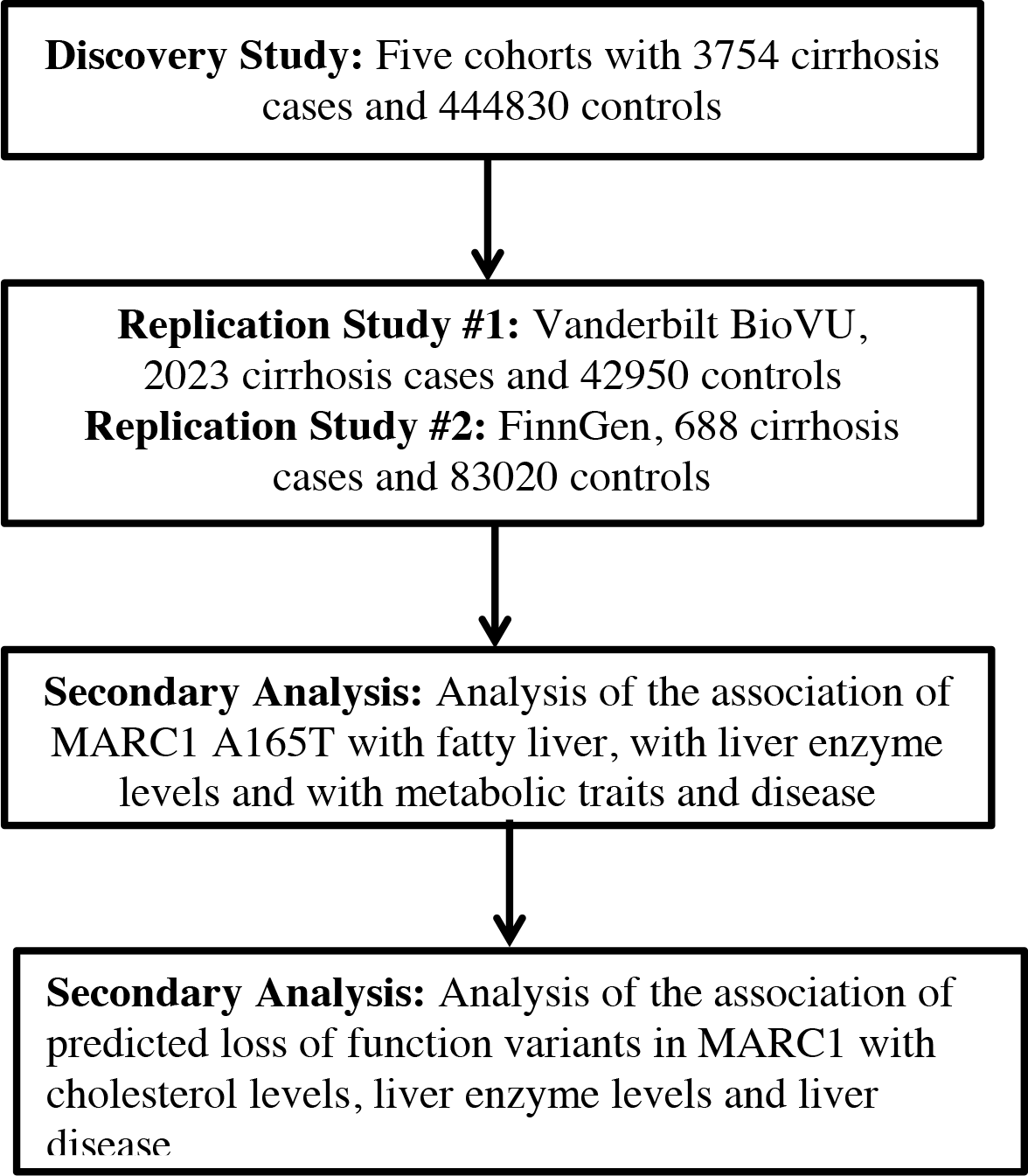
Study design.

To test this hypothesis, we first created an all-cause liver cirrhosis phenotype in UK Biobank, combining the following ICD10 diagnostic codes: K70.2 (alcoholic fibrosis and sclerosis of the liver), K70.3 (alcoholic cirrhosis of the liver), K70.4 (alcoholic hepatic failure), K74.0 (hepatic fibrosis), K74.1 (hepatic sclerosis), K74.2 (hepatic fibrosis with hepatic sclerosis), K74.6 (other and unspecified cirrhosis of liver), K76.6 (portal hypertension), or I85 (esophageal varices). Using this definition, we identified 1740 cases of cirrhosis in UK Biobank. We examined the association of all-cause cirrhosis with six genetic variants previously reported to be associated with alcoholic or non-alcoholic cirrhosis: *PNPLA3* I48M, *TM6SF2* E167K, *MBOAT7* rs641738*, HSD17B13* rs72613567, *HFE* C282Y and *SERPINA1* E366K.^4,9,11,12^ All six variants associated with all-cause cirrhosis in UK Biobank (**Supp. Figure 2**). Each variant exhibited greater statistical significance with all-cause cirrhosis than with alcoholic or non-alcoholic subtypes in UK Biobank, with an average 30% gain in power by analyzing all-cause cirrhosis compared to non-alcoholic cirrhosis and an 87% gain in power compared to analyzing alcoholic cirrhosis.

**Figure 2.**
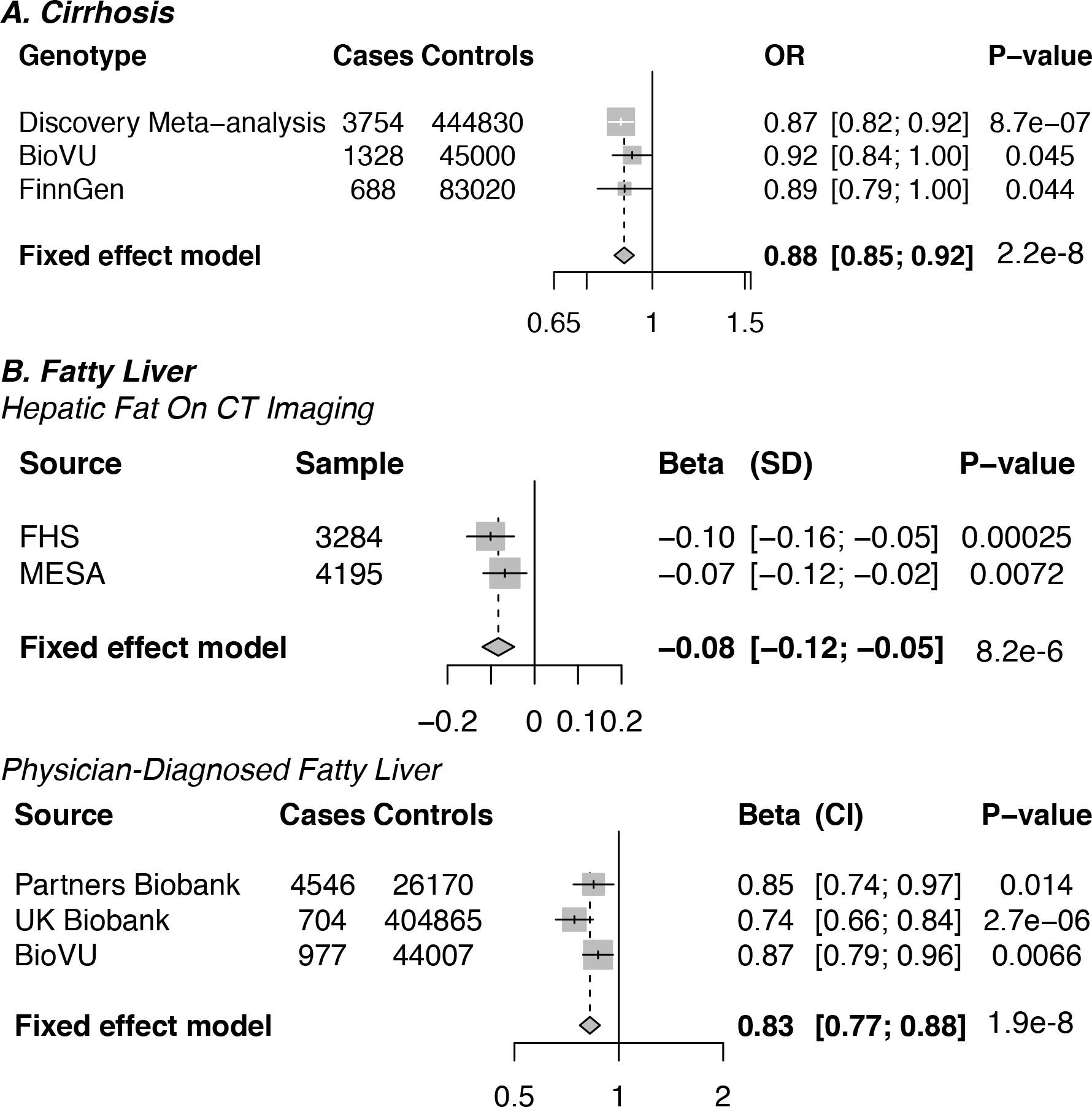
Association of MARC1 p.A165T with cirrhosis and fatty liver in discovery and replication datasets.

Having established that an analysis of all-cause cirrhosis would provide improved statistical power, we sought to identify novel genetic determinants of all-cause cirrhosis through a discovery genome-wide association analysis followed by replication (Figure 1). In the discovery analysis, we analyzed 3,754 all-cause cirrhosis cases and 444,830 controls from five cohorts **(Supp. Table 1)**. We tested the association of 14 million genetic variants with minor allele frequency > 0.1% with all-cause cirrhosis in both additive and recessive models. No evidence of genomic inflation was observed (lambda 1.02, **Supp. Figure 3)**. We replicated three known associations of *PNPLA3*, *TM6SF2* and *HDS17B13* variants with cirrhosis at genome-wide significance (Table 1). We also identified the *HFE* p.C282Y variant (the most cause of hemochromatosis in populations of European ancestry) as associated with all-cause cirrhosis in a recessive model (OR 3.2, p=1.3*10^−14^).^11^ No other variants were associated with all-cause cirrhosis at genome-wide significance.

**Table 1.**
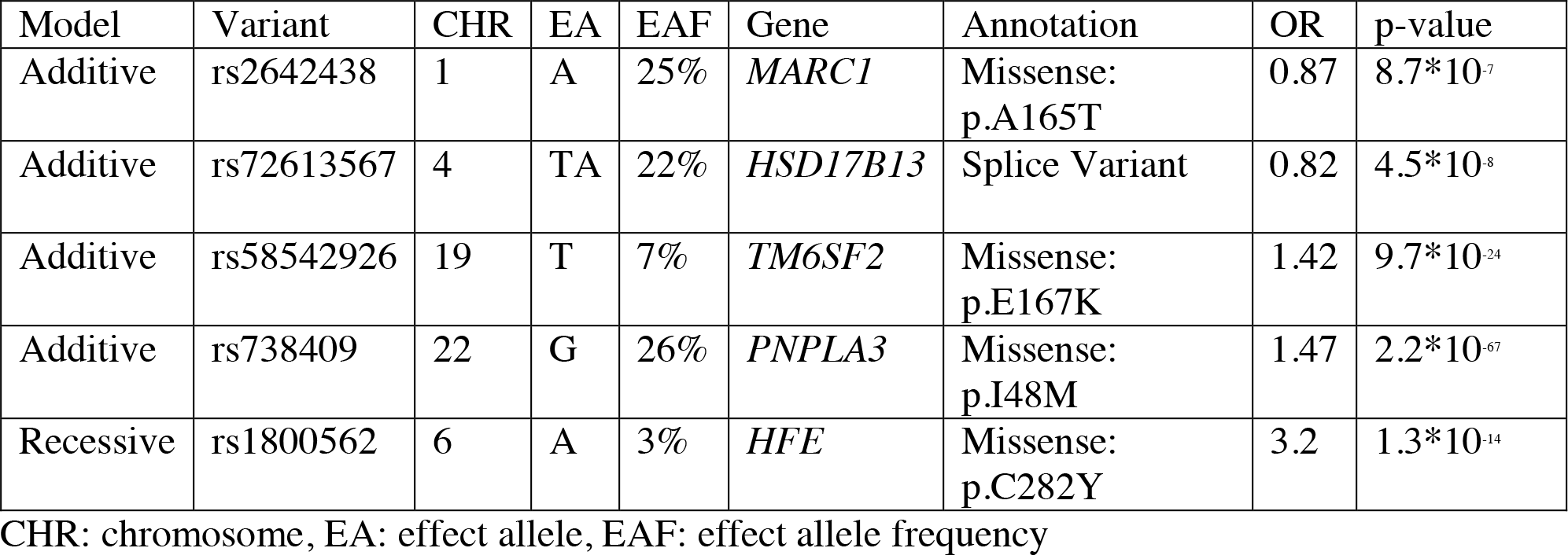
DNA sequence variants associated with all-cause cirrhosis in the discovery analysis.

**Figure 3.**
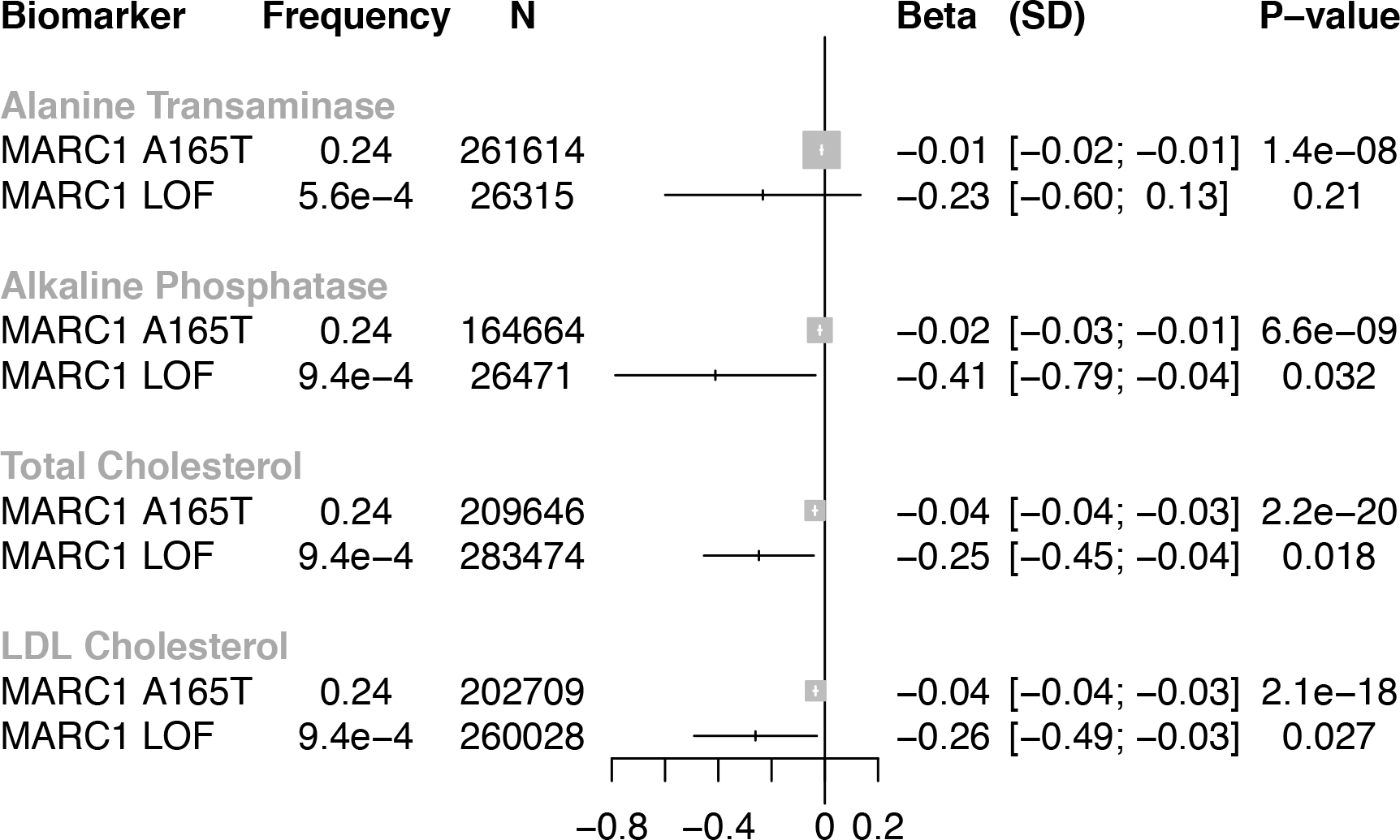
Association of *MARC1* p.A165T and predicted loss of function variants in *MARC1* with alanine transaminase, alkaline phosphatase, total cholesterol and LDL cholesterol.

The lead coding variant at sub-genome wide significance was a missense variant in *MARC1* (p.A165T) that was associated with lower risk of all-cause cirrhosis (OR 0.87, p=8.7*10^−7^, minor allele frequency 25%). We sought replication of this observation in two independent studies – BioVU and the FinnGen Consortium. *MARC1* p.A165T associated with protection from cirrhosis in BioVU (OR 0.92, p=0.045) and in FinnGen (0.89, p=0.044). When the statistical evidence from the discovery and replication studies are combined, *MARC1* p.A165T associated with protection from cirrhosis at a significance level exceeding genome wide significance (OR 0.88, p=2*10^−8^, Figure 2). No evidence of heterogeneity in the association of *MARC1* p.A165T with all-cause cirrhosis in the discovery analysis and replication analyses was observed (p=0.64).

Next, we examined whether *MARC1* p.A165T associated with fatty liver (definitions provided in **Supp. Table 2**). *MARC1* p.A165T was associated with reduced hepatic fat on computed tomographic imaging in both the Framingham Heart Study and Multi-ethnic Study of Atherosclerosis cohorts (−0.08 SD, p=8.2*10^−6^). *MARC1* p.A165T also associated with reduced risk of physician-diagnosed fatty liver in three biobank studies (OR 0.83, p=1.90*10^−8^). In UK Biobank, a similar association of *MARC1* p.A165T with all-cause cirrhosis (OR 0.84, p=1.0*10^−5^) and fatty liver (OR 0.74, p=2.3*10^−6^) was observed when number of medications taken by each participant was adjusted for.

Having established that *MARC1* p.A165T associates with protection from all-cause cirrhosis as well as fatty liver, we tested association of this variant with plasma biomarkers – alanine transaminase (ALT), aspartate transaminase (AST), alkaline phosphatase (ALP), total cholesterol, LDL cholesterol, HDL cholesterol, and triglycerides. *MARC1* p.A165T associated with lower ALT levels (−0.012 SD, p = 1.4*10^−8^) and ALP levels (−0.019 SD, p=6.6*10^−9^) but was not associated with AST levels (−0.005 SD, p=0.12).

In previously published work from the Global Lipids Genetics Consortium, *MARC1* p.A165T associated with lower total cholesterol (−0.037, =1.3*10^−18^) and low-density lipoprotein (LDL) cholesterol (−0.035, p=7.3*10^−16^)^13^. *MARC1* p.A165T associated with higher triglyceride levels (0.017 SD, p=5*10^−6^) and lower HDL levels (−0.030 SD, p=7.8*10^−14^, **Supp. Table 3**) but did not associate with blood pressure, body mass index or waist-to-hip ratio.

Variants in *PNPLA3* and *TM6SF2* that decrease risk of cirrhosis have been reported to increase risk for coronary artery disease (CAD).^14^ This raises the possibility that treatment of cirrhosis (e.g. through *MARC1* or *HSD17B13* inhibition) may have adverse cardiovascular effects. We therefore examined whether *MARC1* p.A165T increases CAD risk. In contrast to *PNPLA3* and *TM6SF2*, neither *MARC1* p.A165T nor *HSD17B13* rs72613567 associated with risk of CAD (**Supp. Figure 4**).^15^ In a phenome wide association study in UK Biobank, *MARC1* p.A165T associated with a lower risk of gallstones (OR 0.96, p=0.0006) and an elevated risk of gout (OR 1.06, p=0.001, **Supp. Figure 5**).

Finally, using two approaches, we addressed whether loss or gain of MARC1 function might be responsible for the protection from cirrhosis and the reduced levels of ALT, ALP, total cholesterol, and LDL cholesterol. In the first approach, we leveraged a rare nonsense mutation observed early in the *MARC1* gene (p.R200Ter). In a combined analysis of UK Biobank, Partners Biobank, MESA and Framingham, one case of liver disease (either fatty liver or cirrhosis) was observed among 110 carriers of *MARC1* R200Ter compared to 7629 cases of liver disease among 446 965 non-carriers (adjusted odds ratio 0.19, p=0.04). Second, we assembled sequence data for *MARC1* in 45,493 individuals and identified 22 carriers of *MARC1* p.R200Ter and 21 carriers of other predicted loss of function variants (**Supp. Tables 4 and 5**). Carriers of predicted loss of function variants in *MARC1* (early truncation, splice site and frameshift variants) had lower ALP levels (−0.41 SD, p=0.032), total cholesterol levels (−0.25 SD, p=0.018) and LDL cholesterol levels (−0.26 SD, p=0.027, Figure 3) and had non-significantly lower ALT levels (−0.23 SD, p=0.21, Figure 3). Combined, these findings indicate that *MARC1* loss of function protects against cirrhosis.

In summary, we identify *MARC1* as a novel genetic determinant of fatty liver and all-cause cirrhosis and suggest that MARC1 deficiency protects against liver disease and reduces blood levels of several biomarkers – ALT, ALP, total cholesterol, and LDL cholesterol. *MARC1* encodes mitochondrial amidoxime-reducing component, a molybdenum-containing enzyme that is involved in metabolizing pro-drugs in the liver.^16^ It contains an N-terminal transmembrane helix that anchors the protein to the outer mitochondrial membrane, with the enzymatic domain of MARC1 located in the cytosol.^17^ The crystal structure of MARC1 was recently described.^18^ The molybdenum cofactor is coordinated in a solvent exposed center by predominantly positively charged amino acids. The A165 residue lies within an alpha helix in the N-terminal domain of MARC1. The function of MARC1 is unknown, however, it has been reported to activate N-hydroxylated prodrugs^19^, reduce nitrite to produce nitric oxide^20^ and detoxify trimethylamine N-oxide.^21^ The mechanism by which *MARC1* may contribute to liver damage and cirrhosis is unclear. The lack of association of *MARC1* p.A165T and *HSD17B1*3 rs72613567 with CAD (in contrast to *PNPLA3* and *TM6SF2*) suggests that pharmacologic treatment of cirrhosis and hepatic steatosis may not universally cause excess cardiovascular risk.

Despite the substantial burden of disease posed by cirrhosis worldwide^22^, identification of genetic risk factors has been limited relative to other common diseases such as type 2 diabetes, coronary artery disease or inflammatory bowel disease. In addition to targeted analyses of cirrhosis subtypes^4^, joint analysis of alcoholic and non-alcoholic cirrhosis cases from multiple cohorts may increase power to identify genetic variants that influence cirrhosis through pathways common to alcoholic and non-alcoholic disease, to identify novel therapeutic targets and to further our understanding of this disease.

## Online Methods

### A. Association of known alcoholic and non-alcoholic cirrhosis variants with all-cause cirrhosis in UK Biobank

To examine whether known alcoholic and non-alcoholic cirrhosis variants associate with all-cause cirrhosis, we tested the association of six known cirrhosis variants (*PNPLA3* I48M, *TM6SF2* E167K, *MBOAT7* rs641738*, HSD17B13* rs72613567, *HFE* C282Y and *SERPINA1* E366K^4,9,11,12^) with all-cause cirrhosis in UK Biobank (ICD codes K70.2, K70.3, K70.4, K74.0, K74.1, K74.2, K74.6, K76.6, or I85). To examine whether this approach increased power relative to examining subtypes of alcoholic and non-alcoholic cirrhosis, we compared the significance of the association of these variants with all-cause cirrhosis (their Z-scores) to the significance of the association of these variants with alcoholic cirrhosis and with non-alcoholic cirrhosis. Alcoholic cirrhosis was defined as physician-diagnosed alcoholic cirrhosis or alcoholic liver failure (ICD codes K70.2, K70.3 or K70.4). Non-alcoholic cirrhosis was defined as non-alcoholic cirrhosis (ICD codes K74.0, K74.1, K74.2, K74.6, K76.6 or I85) that occurred among individuals who drank less than fourteen alcoholic drinks per week. We excluded former drinkers (individuals who previously consumed alcohol but stopped) from analysis of non-alcoholic cirrhosis, as these individuals may have previously consumed alcohol but quit due to adverse effects.^23^ We tested the association of each of the six variants with all-cause cirrhosis, alcoholic cirrhosis and non-alcoholic cirrhosis in UK Biobank using logistic regression adjusted for age, sex, ten principal components of ancestry and a dummy variable for array type.

### B. Genome wide association study for all-cause cirrhosis

We conducted a genome wide association for all-cause cirrhosis using five cohorts: UK Biobank, Partners Biobank, Atherosclerosis Risk in Communities study (ARIC) and summary statistics from two cohorts from a prior genome wide association study of alcoholic cirrhosis.^4^ Definitions of cirrhosis used in each of the five cohorts are provided (**Supp. Table 1**). We excluded cases of cirrhosis secondary to primary biliary cholangitis and primary sclerosis cholangitis as these autoimmune disorders are directed against the biliary (and not hepatic) parenchyma.^24,25^

For UK Biobank, genotyping was performed using either the UK BiLEVE Axiom array or the UK Biobank Axiom array. Phasing and imputation were performed centrally, by UK Biobank, using the Haplotype Reference Consortium and a reference panel of UK 10K merged with the 1000 Genomes phase 3 panel. One related individual of each related pair of individuals, individuals whose genetic sex did not match self-reported sex and individuals with an excess of missing genotype calls or more heterozygosity than expected were excluded from analysis. For Partners Biobank, genotyping was performed using Illumina MEGA array. Variants were imputed to the HapRef consortium using the Michigan Imputation Server.^26^ For ARIC, genotyping was performed using the Affymetrix 6.0 array. Variants were imputed to the HapRef consortium using the Michigan Imputation Server. We excluded any variants with an imputation quality < 0.3.

Genome wide association study in each cohort was performed using logistic regression with adjustment for age, sex and ten principal components of ancestry. We tested the association of fourteen million variants with minor allele frequency of greater than 0.1% with cirrhosis in each cohort. PLINK was used for all analyses. ^27^ To combine estimates across cohorts, inverse variance fixed effects meta-analysis, as implemented by METAL, was used.^28^ Quantile-quantile analysis was used to examine for the presence of population stratification. No evidence of inflation was observed (lambda 1.02; Supplementary Figure 2). Both additive and recessive analyses were performed.

### C. Replication of the association of MARC1 p.A165T with cirrhosis and fatty liver

We replicated the association of *MARC1* p.A165T with all-cause cirrhosis in two cohorts. First, we examined whether *MARC1* p.A165T associates with physician-diagnosed all-cause cirrhosis in the Vanderbilt BioVU, a DNA databank linked to de-identified electronic health records. We identified 46328 individuals of European ancestry with genome-wide genotyping and who were either cases or controls for all-cause cirrhosis. Using ICD-9 (571.2, 571.5, 572.3, 456.0, 456.1, 456.2) or ICD-10 codes (K70.2, K70.3, K70.4, K74.0, K74.1, K74.2, K74.6, K76.6, I85) to define all-cause cirrhosis, 1328 cases were identified. We identified 45000 controls using the EHR-based PheWAS approach, which excludes related diseases based on ICD codes.^29^ Logistic regression, with adjustment for age, sex and principal components of ancestry, was used to estimate the association of *MARC1* p.A165T with cirrhosis in this dataset. Second, in FinnGen, we identified 688 cases of all-cause cirrhosis (ICD-10 K70.2, K70.3, K70.4, K74.0, K74.1, K74.2, K74.6, K76.6, I85) and 83020 controls. Logistic regression, as implemented in SAIGE^30^, was used to test the association of *MARC1* p.A165T with cirrhosis in this dataset while controlling for age, sex and relatedness within the sample.

To examine whether *MARC1* p.A165T associates with fatty liver, we tested the association of this variant with fatty liver in five cohorts. In the Framingham cohort (Offspring Cohort and Third Generation Cohort), we examined whether MARC1 p.A165T associates with hepatic steatosis on CT imaging. 3284 individuals in Framingham with genotype data available underwent multidetector abdominal CT.^31^ We measured hepatic steatosis by computing the liver-to-phantom ratio of the average Hounsfield units of three liver measurements to average Hounsfield units of three phantom measurements (to correct for inter-individual differences in penetration), as previously described.^31^ We tested the association of the p.A165T variant with liver-to-phantom ratio with adjustment for age, sex and ten principal components of ancestry using a linear mixed model to control for relatedness among individuals. In the Multi-Ethnic Study of Atherosclerosis cohort (MESA), 4195 individuals underwent multidetector CT. Hepatic steatosis was measured as the mean of three attenuation measurements (two in the right lobe of the liver and one in the left lobe). No phantom measurement was available for standardization. We therefore tested the association of the p.A165T variant with mean liver attenuation with adjustment for age, sex and ten principal components of ancestry. Individuals with higher liver fat have *lower* liver-to-phantom ratios and liver attenuation measurements. For interpretability, we therefore report all estimates in units of standard deviation increases in liver fat, with a one standard deviation increase in liver fat corresponding to a one standard deviation decrease in the liver-to-phantom ratio or mean liver attenuation.

In three cohorts (Partners Biobank, UK Biobank, BioVU), we lacked CT imaging data to measure hepatic steatosis. We therefore tested the association of the *MARC1* p.A165T variant with physician-diagnosed fatty liver in these cohorts (ICD codes K76.0 fatty change of liver, K76.5 non-alcoholic steatohepatitis) using logistic regression, adjusted for age, sex and ten principal components of ancestry. We pooled estimates of the association of *MARC1* p.A165T with fatty liver across all five cohorts using fixed effects meta-analysis.^28^

### D. Association of MARC1 p.A165T with liver enzyme levels, metabolic traits and disease

We tested the association of p.A165T with metabolic traits using four different datasets. For serum levels of liver enzymes, we used data from Partners Biobank (n=26471), Framingham (n=3288), LOLIPOP (n=54857)^32^ and BioBank Japan (n=134182)^33^ where measures of serum alanine transaminase (ALT), aspartate transaminase (AST) and alkaline phosphatase (ALP) were available. We log transformed ALT, AST and ALP. We then conducted a linear regression analysis with adjustment for age, sex and ten principal components of ancestry. We pooled estimates across cohorts using inverse variance weighted fixed effects meta-analysis. For lipids (LDL cholesterol, HDL cholesterol, triglycerides and total cholesterol), we used data from the Global Lipids Genetics Consortium, a meta-analysis of 188 587 individuals of European descent.^13^ This GWAS included 37 studies genotyped using the Illumina Metabochip array as well as an additional 23 studies genotyped using a variety of arrays. For BMI and WHRadjBMI we used data from the Genetic Investigation of ANthropometric Traits (GIANT) consortium.^34,35^ For WHRadjBMI, data from 210,088 individuals of European ancestry were included. For BMI, data for 322,154 individuals of European ancestry were included. Individuals were genotyped using various arrays and imputed with the HapMap reference panel to 2.5 million SNPs. For blood pressure, we used data from UK Biobank. We tested the association of p.A165T with systolic blood pressure and diastolic blood pressure using linear regression with adjustment for age, sex and ten principal components of ancestry.

### E. Phenome-wide association study for MARC1 p.A165T in UK Biobank

A phenome-wide association study of *MARC1* p.A165T in UK Biobank was performed. ^36,37^ We tested the association of MARC1 p.A165T with diseases with more than one thousand cases in UK Biobank. Definitions for 31 different diseases analyzed in the phenome-wide association study are provided (**Supp. Table 6**). The association of p.A165T with each disease was estimated using logistic regression with adjustment for age, sex, ten principal components of ancestry and a dummy variable for array type. A Bonferroni adjusted significance level of p < 0.0016 (0.05/31) was used.

### F. Association of rare predicted loss of function variants in MARC1 with cholesterol levels, liver enzyme levels and liver disease

*MARC1* p.A165T associated with reduced total cholesterol levels, LDL cholesterol levels, alanine transaminase levels and alkaline phosphatase levels. To examine whether *MARC1* deficiency may therefore protect against elevated cholesterol levels and liver enzyme levels, we estimated the association of rare predicted loss of function variants in *MARC1* with these phenotypes, using five data sources. First, we examined the association of a rare genotyped stop codon in *MARC1* (Arg200Ter, rs139321832) with total and LDL cholesterol in the Global Lipids Genetics Consortium exome chip analysis of 283474 individuals.^14^ Second, we examined the association of rare predicted loss of function variants with cholesterol in two exome sequence datasets: the Myocardial Infarction Genetics consortium (n=27034) and the T2D Genes consortium (n=18456). Sequence data for *MARC1* were extracted from exome sequencing performed in the MIGen Consortium as previously described.^38,39^ The Burrows–Wheeler Aligner algorithm was used to align reads from participants to the reference genome (hg19). The GATK HaploTypeCaller was used to jointly call variants. Metrics including Variant Quality Score Recalibration (VQSR), quality over depth, and strand bias were then used to filter variants. The Jackson Heart Study was excluded from analysis of MIGen as it was included in the T2D Genes consortium. Exome sequencing was performed in the T2D Genes consortium as previously described.^40^ To analyze exome sequences from the T2D Genes consortium, the online Genetic Association Interactive Test in the T2D Knowledge portal was used.^40^

Predicted loss of function variants in *MARC1* were defined as those which resulted in loss-of-function of the protein (nonsense mutations that resulted in early termination of *MARC1*, frameshift mutations due to insertions or deletions of DNA, or splice-site mutations which result in an incorrectly spliced protein), as previously described.^41^ The Variant Effect Predictor algorithm was used to annotate predicted damaging variants.^42^ We excluded the variant Arg200Ter from analysis in MIGEN and T2D Genes to prevent overlap between samples. Estimates were adjusted for age, sex and five principle components of ancestry. We pooled estimates from all three data sources using inverse variance weighted fixed effects meta-analysis.

Third, we examined the association of the genotyped Arg200Ter variant with log-transformed ALT and ALP levels using data from the Partners Biobank cohort. We tested for the association of this variant with log-transformed ALT and ALP levels using linear regression with adjustment for age, sex and ten principal components of ancestry.

## Acknowledgements

This research has been conducted using the UK Biobank resource, application 7089. This work was funded by the National Institutes of Health (R01 HL127564 to S.K.), which had no involvement in the design and conduct of the study; the collection, analysis, and interpretation of the data; or the preparation, review, and approval of the manuscript. Samples for the Leicester cohort were collected as part of projects funded by the British Heart Foundation (British Heart Foundation Family Heart Study, RG2000010; UK Aneurysm Growth Study, CS/14/2/30841) and the National Institute for Health Research (NIHR Leicester Cardiovascular Biomedical Research Unit Biomedical Research Informatics Centre for Cardiovascular Science, IS_BRU_0211_20033). NJS is supported by the British Heart Foundation and is a NIHR Senior Investigator. The Munich MI Study is supported by the German Federal Ministry of Education and Research (BMBF) in the context of the e:Med program (e:AtheroSysMed) and the FP7 European Union project CVgenes@target (261123). Additional grants were received from the Fondation Leducq (CADgenomics: Understanding Coronary Artery Disease Genes, 12CVD02).This study was also supported through the Deutsche Forschungsgemeinschaft cluster of excellence “Inflammation at Interfaces” and SFB 1123. The Italian Atherosclerosis, Thrombosis, and Vascular Biology (ATVB) Study was supported by a grant from RFPS-2007-3-644382 and Programma di ricerca Regione-Università 2010-2012 Area 1-Strategic Programmes-Regione Emilia-Romagna. Funding for the exome-sequencing project (ESP) was provided by RC2 HL103010 (HeartGO), RC2 HL102923 (LungGO), and RC2 HL102924 (WHISP). Exome sequencing was performed through RC2 HL102925 (BroadGO) and RC2 HL102926 (SeattleGO). The JHS is supported by contracts HHSN268201300046C, HHSN268201300047C, HHSN268201300048C, HHSN268201300049C, HHSN268201300050C from the National Heart, Lung, and Blood Institute and the National Institute on Minority Health and Health Disparities. Dr. Wilson is supported by U54GM115428 from the National Institute of General Medical Sciences. Exome sequencing in ATVB, PROCARDIS, Ottawa, PROMIS, Southern German Myocardial Infarction Study, and the Jackson Heart Study was supported by 5U54HG003067 (to Dr. Gabriel).

## Author Contributions

Concept and design: C.A.E., M.H., S.K. Acquisition, analysis, or interpretation of data: C.A.E., S.K. Drafting of the manuscript: C.A.E., M.H., S.K. Critical revision of the manuscript for important intellectual content: All authors. Administrative, technical, or material support: S.K.

