## Supplementary Appendix for "A missense variant in Mitochondrial Amidoxime Reducing Component 1 gene and protection against liver disease"

**Online Supplement**

Supplementary Table 1. Definition of cirrhosis in each cohort.

| Cohort | Definition of cirrhosis | Cases | Controls | Individual-level data |
|---|---|---|---|---|
| UK Biobank | Hospitalization or death due to physician diagnosed cirrhosis: K70.2 (alcoholic fibrosis and sclerosis of the liver), K70.3 (alcoholic cirrhosis), K70.4 (alcoholic hepatic failure), K74.0 (hepatic fibrosis), K74.1 (hepatic sclerosis), K74.2 (hepatic fibrosis with hepatic sclerosis), K74.6 (other and unspecific cirrhosis of liver), K76.6 (portal hypertension), or I85 (esophageal varices) | 1740 | 403829 | Yes |
| Partners Biobank | Hospitalization or death due to physician diagnosed cirrhosis: K70.2 (alcoholic fibrosis and sclerosis of the liver), K70.3 (alcoholic cirrhosis), K70.4 (alcoholic hepatic failure), K74.0 (hepatic fibrosis), K74.1 (hepatic sclerosis), K74.2 (hepatic fibrosis with hepatic sclerosis), K74.6 (other and unspecific cirrhosis of liver), K76.6 (portal hypertension), or I85 (esophageal varices) | 1214 | 29502 | Yes |
| ARIC | Hospitalization or death due to physician diagnosed cirrhosis: K70.2 (alcoholic fibrosis and sclerosis of the liver), K70.3 (alcoholic cirrhosis), K70.4 (alcoholic hepatic failure), K74.0 (hepatic fibrosis), K74.1 (hepatic sclerosis), K74.2 (hepatic fibrosis with hepatic sclerosis), K74.6 (other and unspecific cirrhosis of liver), K76.6 (portal hypertension), or I85 (esophageal varices) | 88 | 10034 | Yes |
| Alcoholic cirrhosis GWAS: German Cohort | Presence of cirrhosis on liver biopsy (fibrosis stage 5 or 6) or unequivocal clinical and laboratory evidence for the presence of cirrhosis | 410 | 1119 | No |
| Alcoholic cirrhosis GWAS: UK Cohort | Histological examination of liver tissue or compatible historical, clinical, laboratory, radiological and endoscopic features | 302 | 346 | No |
| Total | | 3754 | 444830 | |

Supplementary Table 2. Definition of fatty liver in each cohort.

| Cohort | Definition of cirrhosis | Cases | Controls | Individual-level data |
|---|---|---|---|---|
| Framingham | CT: Ratio of mean of liver attenuation measurements to phantom measurement | 3284 | | Yes |
| MESA | CT: Mean of three liver attenuation measurements | 4195 | | Yes |
| UK Biobank | Physician diagnosed: K76.0 (fatty change of liver), K76.5 (non-alcoholic steatohepatitis) | 704 | 404865 | Yes |
| Partners Biobank | Physician diagnosed: K76.0 (fatty change of liver), K76.5 (non-alcoholic steatohepatitis) | 4546 | 26170 | Yes |
| BioVU | Physician diagnosed: K76.0 (fatty change of liver), K76.5 (non-alcoholic steatohepatitis) | 977 | 44007 | Yes |
| Total | | 488748 | | |

Supplementary Table 3. Association of MARC1 A165T with metabolic traits.

| ***Outcome*** | ***Source*** | ***n*** | ***Beta*** | ***SE*** | ***p-value*** |
|---|---|---|---|---|---|
| *Liver enzymes* | | | | | |
| ALT | Partners Biobank, Framingham, LOLIPOP, BioBank Japan, BioVU | 261614 | -0.012 | 0.003 | 1.4*10^-8 |
| AST | Partners Biobank, Framingham, LOLIPOP, BioBank Japan | 221662 | -0.005 | 0.003 | 0.12 |
| AP | Partners Biobank, Framingham, LOLIPOP, BioBank Japan | 164664 | -0.019 | 0.003 | 6.6*10^-9 |
| *Blood lipids* | | | | | |
| Total Cholesterol | GLGC | 188577 | -0.037 SD | 0.004 | 1.3*10^-18 |
| LDL Cholesterol | GLGC | 188577 | -0.035 SD | 0.004 | 7.3*10^-16 |
| HDL Cholesterol | GLGC | 188577 | -0.030 SD | 0.004 | 7.8*10^-14 |
| Triglycerides | GLGC | 188577 | 0.017 SD | 0.004 | 5.3*10^-6 |
| *Blood pressure* | | | | | |
| Systolic blood pressure | UK Biobank | 379771 | 0.001 SD | 0.002 | 0.55 |
| Diastolic blood pressure | UK Biobank | 379782 | 0.003 SD | 0.003 | 0.19 |
| *Anthropometric measurements* | | | | | |
| Body mass index | GIANT | 325053 | 0.005 SD | 0.003 | 0.17 |
| Waist to hip ratio adjusted for body mass index | GIANT | 211816 | 0.006 SD | 0.004 | 0.15 |

GLGC: Global lipids genetics consortium, GIANT: Genetic investigation of anthropometric traits consortium, SD: standard deviations

Supplementary Table 4. Rare predicted loss of function variants in MIGEN

| CHR:POS_REF/ALT | Consequence | Amino Acid Change | Individuals With Variant |
|---|---|---|---|
| 1:220970148_G/A | Splice Donor | | 2 |
| 1:220970148_G/C | Splice Donor | | 1 |
| 1:220971357_G/A | Splice Donor | | 1 |
| 1:220978456_G/A | Splice Donor | | 2 |
| 1:220978457_T/C | Splice Donor | | 1 |
| 1:220986659_C/T | Stop Gained | Arg305Ter | 2 |
| 1:220986755_C/T | Stop Gained | Gln337Ter | 3 |
| Total | | | 12 |

Supplementary Table 5. Rare predicted loss of function variants in T2D Genes.

| CHR:POS_REF/ALT | Consequence | Amino Acid Change | Individuals With Variant |
|---|---|---|---|
| 1:220960469_TG/T | Frameshift | Trp62fs | 2 |
| 1:220960562_G/A | Splice Donor | | 1 |
| 1:220970097_C/T | Stop Gained | Arg188Ter | 1 |
| 1:220978402_G/A | Stop Gained | Trp254Ter | 2 |
| 1:220986659_C/T | Stop Gained | Arg305Ter | 2 |
| 1:220986728_TG/T | Frameshift | Val328fs | 1 |
| Total | | | 9 |

Supplementary Table 6. Definition of outcomes in phenome wide association study in UK Biobank.

| Outcome | Definition (UK Biobank unless otherwise specified) |
|---|---|
| Coronary artery disease | (1) Myocardial infarction (MI), coronary artery bypass grafting, or coronary artery angioplasty documented in medical history at time of enrollment by a trained nurse or<br>(2) Hospitalization for ICD-10 code for acute myocardial infarction (I21.0, I21.1, I21.2, I21.4, I21.9) or<br>(3) Hospitalization for OPCS-4 coded procedure: coronary artery bypass grafting (K40.1-40.4, K41.1-41.4, K45.1-45.5) or<br>(4) Hospitalization for OPCS-4 coded procedure: coronary angioplasty ± stenting (K49.1-49.2, K49.8-49.9, K50.2, K75.1-75.4, K75.8-75.9) |
| Atrial fibrillation/flutter | History of atrial fibrillation or flutter during verbal interview with trained nurse or hospitalization for or death due to ICD code I48 |
| Heart failure | History of heart failure during verbal interview with trained nurse or hospitalization for or death due to ICD code I11.0, I13.0, I13.2, I125.5, I42, I50 |
| Stroke | History of stroke, adjudicated by UK Biobank centrally as report of stroke during verbal interview with trained nurse or hospitalization for or death due to ICD code I60-64 *(http://biobank.ctsu.ox.ac.uk/crystal/refer.cgi?id=462)* |
| Peripheral vascular disease | History of peripheral vascular disease or intermittent claudication during verbal interview with trained nurse or hospitalization for or death due to ICD code I70, I73.8 or I73.9 |
| Venous thromboembolism | History of venous thromboembolism, deep vein thrombosis or pulmonary embolism during verbal interview with trained nurse or hospitalization for death due to I26, I80.1, I80.2, I81, or I82.0 |
| Aortic stenosis | History of aortic stenosis during verbal interview with trained nurse or hospitalization for ICD code I06.0, I06.2 I35.0 or I35.2 |
| Inflammatory bowel disease | History of inflammatory bowel disease, Crohn's disease or ulcerative colitis during verbal interview with trained nurse or hospitalization for or death due to ICD code K50 or K51 |
| Gastric reflux | History of gastric reflux during verbal interview with trained nurse or hospitalization for or death due to ICD code K21 |
| Gallstone | History of gallstones during verbal interview with trained nurse or hospitalization for or death due to ICD code K56.3 or K80 |
| Type 2 Diabetes | History of diabetes unspecified, type 2 diabetes during verbal interview with trained nurse or hospitalization for or death due to ICD code E11 |
| Hyperthyroidism | History of hyperthyroidism during verbal interview with trained nurse or hospitalization for or death due to ICD code E05 |
| Hypothyroidism | History of hypothyroidism during verbal interview with trained nurse or hospitalization for or death due to ICD code E03 |
| Gout | History of gout during verbal interview with trained nurse or hospitalization for or death due to ICD code M10 |
| Enlarged prostate | History of enlarged prostate during verbal interview with trained nurse or hospitalization for or death due to ICD code N40 |
| Uterine fibroids | History of uterine fibroids during verbal interview with trained nurse or hospitalization for or death due to ICD code D25 |
| Migraine | History of migraine during verbal interview with trained nurse or hospitalization for or death due to ICD code G43 |
| Depression | History of depression during verbal interview with trained nurse or hospitalization for or death due to ICD code F32 |
| Anxiety | History of anxiety/panic attacks during verbal interview with trained nurse or hospitalization for or death due to ICD code F41 |

| | |
|---|---|
| Osteoporosis | History of osteoporosis during verbal interview with trained nurse or hospitalization for or death due to ICD code M80 or M81 |
| Osteoarthritis | History of osteoarthritis during verbal interview with trained nurse or hospitalization for or death due to ICD code M15-19 |
| Sciatica | History of sciatica during verbal interview with trained nurse or hospitalization for or death due to ICD code M54.3 |
| Prolapsed disc | History of prolapsed disc/slipped disc during verbal interview with trained nurse or hospitalization for or death due to ICD code M50.2 or M51.2 |
| Asthma | History of asthma during verbal interview with trained nurse or hospitalization for or death due to ICD code J45 or J46 |
| COPD/Emphysema | History of chronic obstructive airways disease, emphysema/chronic bronchitis or emphysema during verbal interview with trained nurse or hospitalization for or death due to ICD code J41-44 |
| Pneumonia | History of pneumonia during verbal interview with trained nurse or hospitalization for or death due to ICD code J12-18 |
| Hayfever | History of hayfever during verbal interview with trained nurse or hospitalization for or death due to ICD code J30 |
| Lung cancer | History of lung cancer during verbal interview with trained nurse or hospitalization for or death due to ICD code C34 |
| Colorectal cancer | History of large bowel cancer/colorectal cancer, colon cancer/sigmoid cancer or rectal cancer during verbal interview with trained nurse or hospitalization for or death due to ICD code C18 |
| Skin cancer | History of skin cancer, malignant melanoma, non-melanoma skin cnacer, basal cell carcinoma or squamous cell carcinoma during verbal interview with trained nurse or hospitalization for or death due to ICD code C43-44 |
| Prostate cancer | History of prostate cancer during verbal interview with trained nurse or hospitalization for or death due to ICD code C61 |
| Cervical cancer | History of cervical cancer or cin cells at the cervix during verbal interview with trained nurse or hospitalization for or death due to ICD code C53 |

Abbreviations: COPD, chronic obstructive pulmonary disease; ICD, international classification of disease

| Type of Cirrhosis | Source | Cases | Controls | OR | 95% CI | p−value |
|---|---|---|---|---|---|---|
| **PNPLA3 I48M** | | | | | | |
| Alcoholic Cirrhosis | Buch et al. [1] | 1860 | 2348 | 2.39 | [2.05; 2.78] | 1.5e−48 |
| Non−Alcoholic Cirrhosis | Speliotes et al. [2] | 533 | 1405 | 3.37 | [2.86; 3.98] | 3.6e−46 |
| Hepatitis C Cirrhosis | Valenti et al. [3] | 215 | 604 | 1.47 | [1.15; 1.87] | 0.002 |
| **TM6SF2 E167K** | | | | | | |
| Alcoholic Cirrhosis | Buch et al. [1] | 1860 | 2348 | 1.84 | [1.42; 2.39] | 7.9e−10 |
| Non−Alcoholic Cirrhosis | Liu et al. [4] | 437 | 637 | 1.88 | [1.41; 2.50] | 1.6e−05 |
| Hepatitis C Cirrhosis | Liu et al. [5] | . | . | 2.05 | [1.39; 3.02] | 3e−04 |
| **HSD17B13 rs72613567:TA** | | | | | | |
| Alcoholic Cirrhosis | Abul−Husn et al. [6] | 124 | 29928 | 0.56 | [0.41; 0.77] | 0.00034 |
| Non−Alcoholic Cirrhosis | Abul−Husn et al. [6] | 374 | 29928 | 0.74 | [0.62; 0.88] | 0.00048 |
| Hepatitis C Cirrhosis | About et al. [7] | 40 | 48 | 0.38 | [0.16; 0.91] | 0.03 |

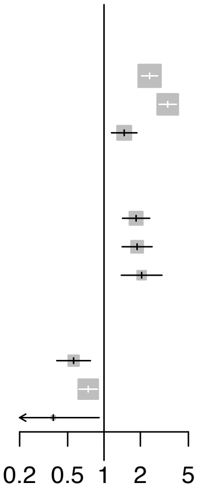


Supplementary Figure 1. Risk of alcoholic, non-alcoholic and hepatitis C cirrhosis associated with PNPLA3 I48M, TM6SF2 E40K and HSD17B13.[1-7] Valenti et al. reports results from a recessive model.

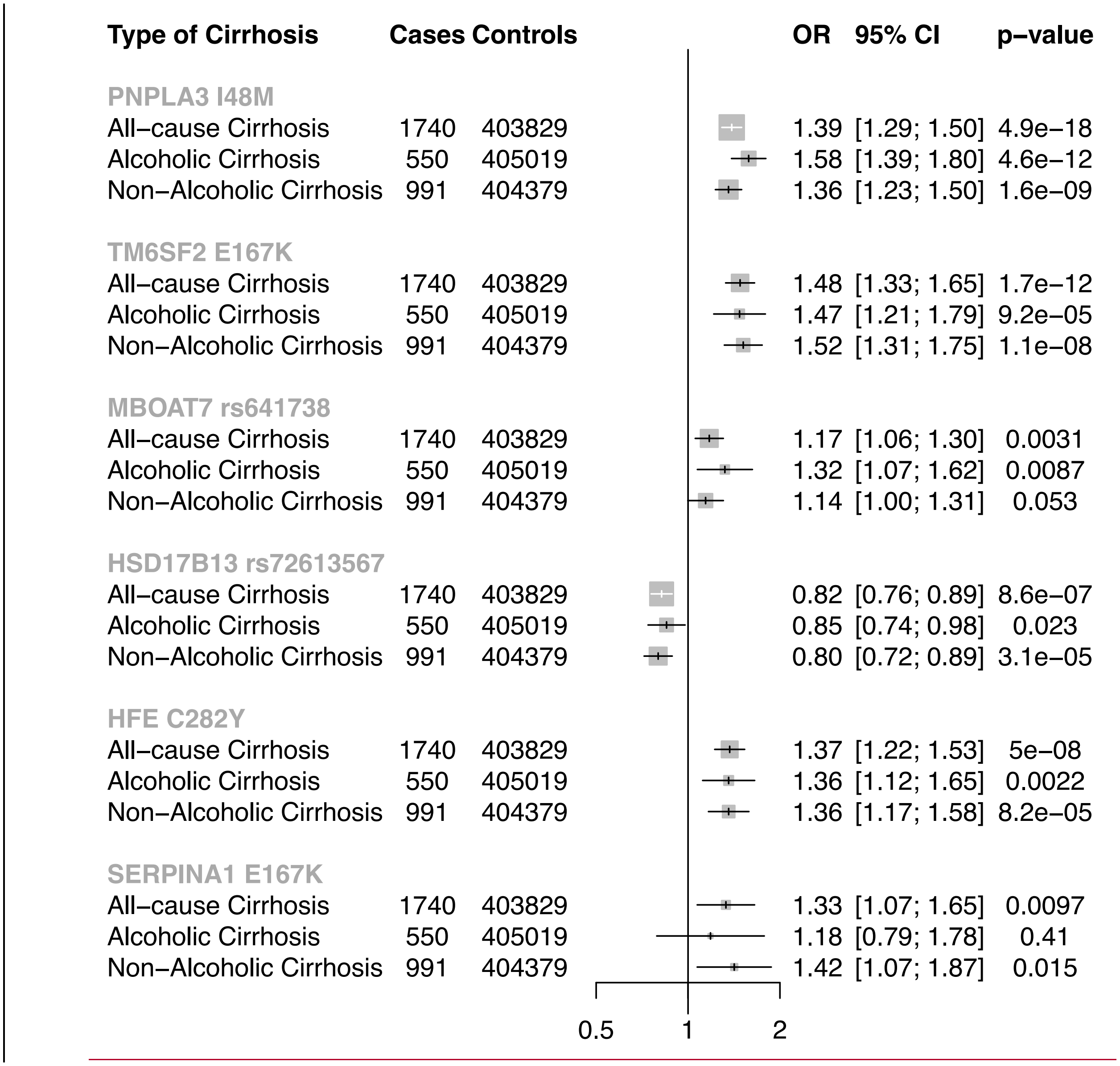


Supplementary Figure 2. Association of known alcoholic and non-alcoholic cirrhosis variants with all-cause cirrhosis in UK Biobank.

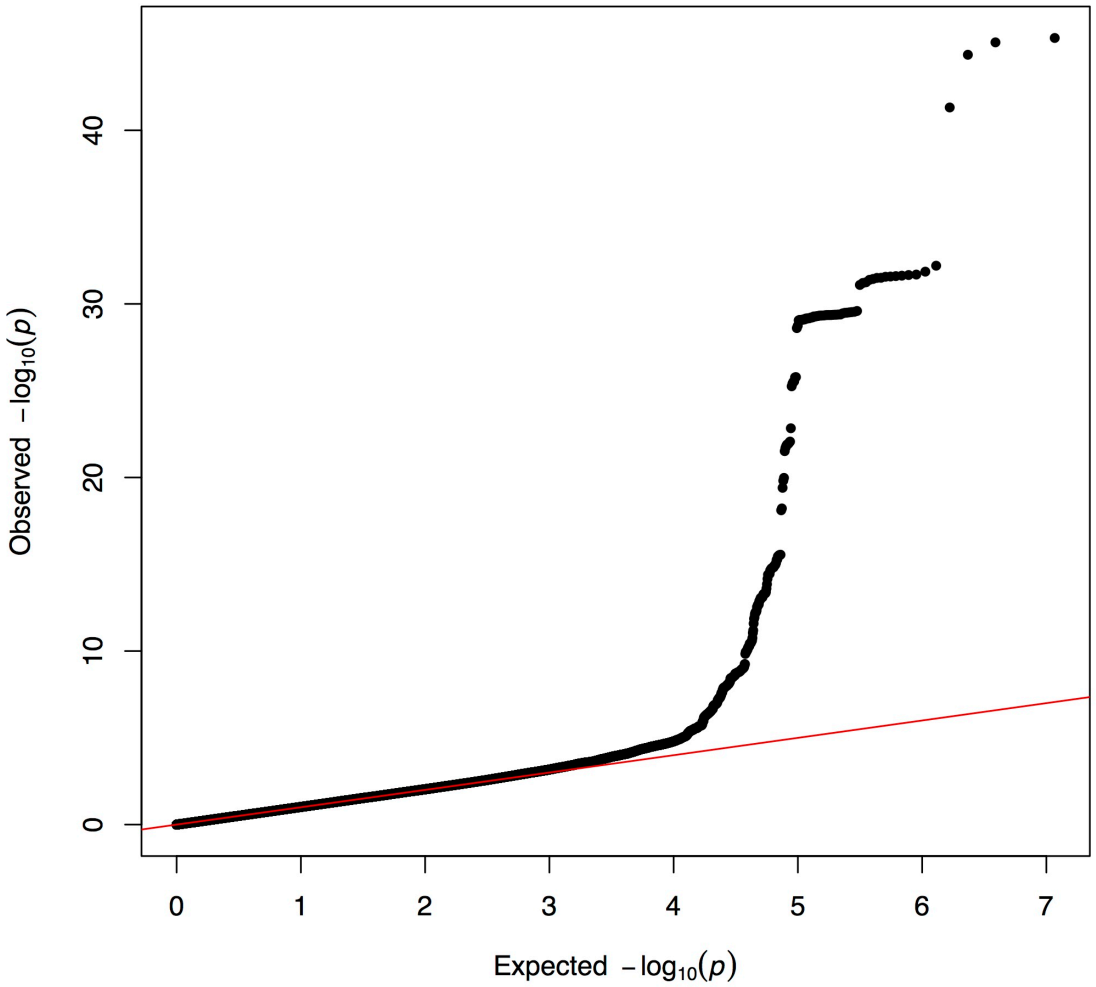


Supplementary Figure 3. QQ plot for genome wide analysis of cirrhosis. Lambda = 1.02

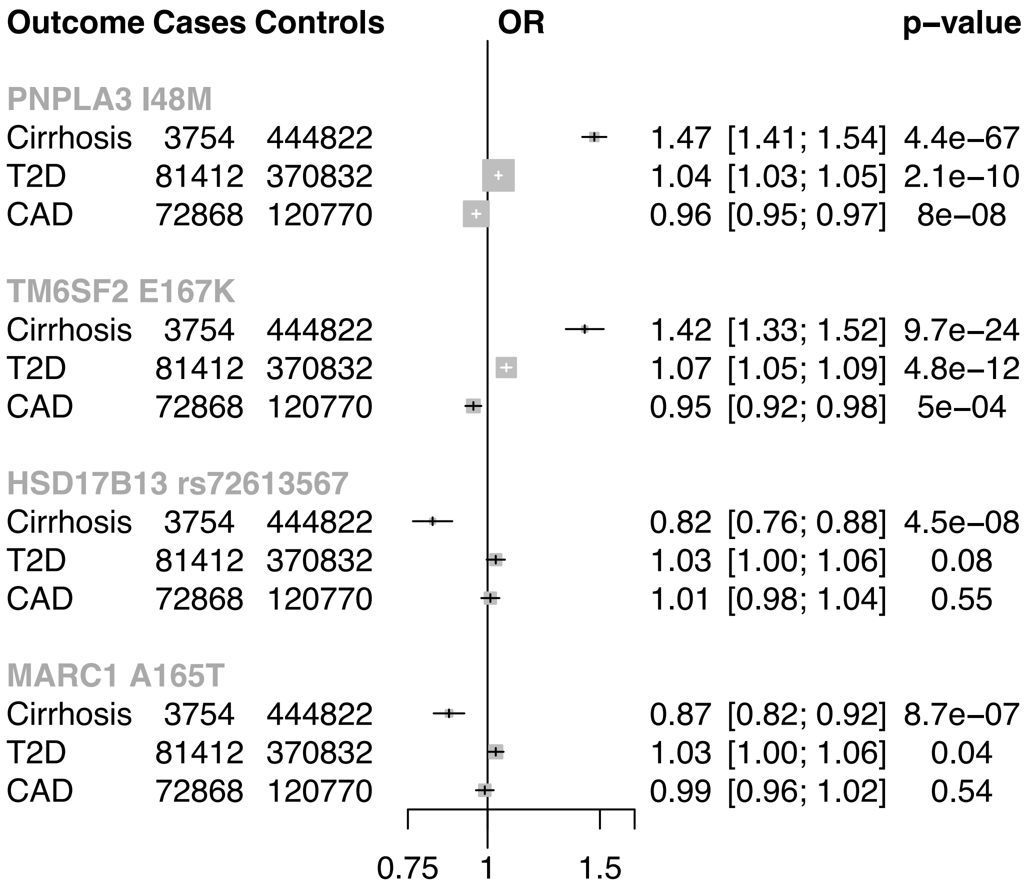


Supplementary Figure 4. Association of cirrhosis variants with type 2 diabetes, coronary artery disease and cirrhosis.

| Outcome | Cases | Controls | OR | | p–value |
|---|---|---|---|---|---|
| **Cardiovascular** | | | | | |
| Coronary Artery Disease | 14811 | 390758 | 1.01 | [0.98; 1.04] | 0.49 |
| Stroke | 9227 | 396342 | 1.00 | [0.97; 1.04] | 0.85 |
| Heart Failure | 6039 | 399530 | 1.02 | [0.98; 1.06] | 0.31 |
| Atrial Fibrillation | 14623 | 390946 | 1.01 | [0.98; 1.04] | 0.47 |
| Aortic Stenosis | 1850 | 403719 | 0.96 | [0.89; 1.03] | 0.26 |
| Peripheral vascular disease | 4720 | 400849 | 1.03 | [0.98; 1.08] | 0.21 |
| Venous thromboembolism | 13281 | 392288 | 1.00 | [0.97; 1.03] | 0.87 |
| **Gastrointestinal disease** | | | | | |
| Inflammatory bowel disease | 5400 | 400169 | 0.96 | [0.92; 1.01] | 0.092 |
| Gastric reflux | 35568 | 370001 | 1.00 | [0.99; 1.02] | 0.82 |
| Gallstones | 17780 | 387789 | 0.96 | [0.94; 0.98] | 0.00064 |
| **Endocrine** | | | | | |
| Type 2 Diabetes | 20458 | 385111 | 1.00 | [0.98; 1.02] | 0.85 |
| Hyperthyroidiusm | 4477 | 401092 | 1.01 | [0.97; 1.06] | 0.65 |
| Hypothyroidism | 23591 | 381978 | 0.99 | [0.97; 1.01] | 0.55 |
| Gout | 7382 | 398187 | 1.06 | [1.02; 1.10] | 0.0013 |
| **Urological** | | | | | |
| Enlarged Prostate | 14323 | 391246 | 0.99 | [0.96; 1.01] | 0.3 |
| Uterine Fibroids | 17128 | 388441 | 0.98 | [0.96; 1.01] | 0.17 |
| **Neurological/psychiatric** | | | | | |
| Migraine | 14293 | 391276 | 1.00 | [0.97; 1.02] | 0.84 |
| Depression | 30835 | 374734 | 1.02 | [1.00; 1.04] | 0.07 |
| Anxiety | 6733 | 398836 | 1.02 | [0.98; 1.06] | 0.3 |
| **Muscoskeletal** | | | | | |
| Osteoporosis | 10863 | 394706 | 0.98 | [0.95; 1.01] | 0.29 |
| Osteoarthritis | 62643 | 342926 | 1.02 | [1.00; 1.03] | 0.01 |
| Sciatica | 6222 | 399347 | 0.98 | [0.95; 1.02] | 0.44 |
| Prolapsed disc | 9463 | 396106 | 1.01 | [0.98; 1.05] | 0.39 |
| **Respiratory** | | | | | |
| Asthma | 47179 | 358390 | 1.01 | [0.99; 1.02] | 0.49 |
| COPD/Emphysema | 13771 | 391798 | 0.99 | [0.97; 1.02] | 0.6 |
| Pneumonia | 14869 | 390700 | 0.99 | [0.96; 1.01] | 0.34 |
| Hayfever | 25489 | 380080 | 1.00 | [0.98; 1.02] | 0.66 |
| **Cancer** | | | | | |
| Lung Cancer | 2281 | 403288 | 1.02 | [0.96; 1.09] | 0.57 |
| Colorectal Cancer | 3969 | 401600 | 0.99 | [0.95; 1.04] | 0.78 |
| Skin Cancer | 16904 | 388665 | 0.99 | [0.97; 1.02] | 0.65 |
| Prostate Cancer | 6377 | 399192 | 1.04 | [1.00; 1.08] | 0.046 |
| Cervical Cancer | 1707 | 403862 | 1.05 | [0.98; 1.13] | 0.17 |

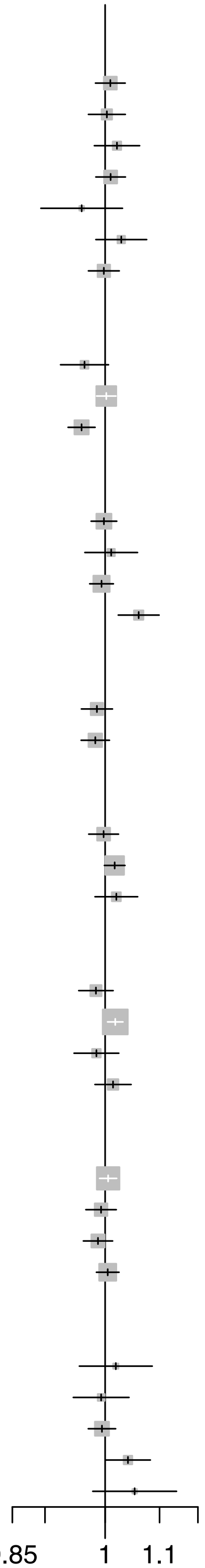


Supplementary Figure 5. Association of MARC A165T with other diseases in a phenome wide association study.
